# A fractional order network model for ZIKA

**DOI:** 10.1101/039917

**Authors:** H. Elsaka, E. Ahmed

## Abstract

Zika is a fast spreading epidemic. So far it is known to have two transmission routes one via mosquito and the other is via sexual contact. It is dangerous on pregnant women otherwise it is mild

## 1 Introduction

Zika is a fast spreading epidemic. Within one year it has spread to more than 32 countries including south, central and north America. One case appeared in Europe and in Australia. Therefore it should be studied using networks [22]. So far it is known to have two transmission routes one via mosquito and the other is via sexual contact. It is dangerous on pregnant women otherwise it is mild or asymptomatic. Therefore we expect that in the future the second route will be more difficult to control. Fractional order (FO) models [14-18] are quite useful in epidemic models to predict the spread of diseases, how to prevent epidemics and so much more. Therefore we present a fractional order network model for ZIKA. The benefit of simple models is that we can average out some of this complexity and try to understand the big picture. Our model will be useful as a conceptual tool for modeling the impact of interventions aiming to control the disease.

In sec.2 a brief introduction to FO calculus is given. In sec.3 the model is given and studied. Sec.4 contains our conclusions.

## 2 Fractional order calculus

### Definition 1

The fractional integral of order *β* ϵ *R*^+^ of the function *f*(*t*), *t* > 0 is defined by

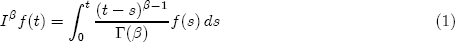

and the fractional derivative of order *α* ϵ (*n* − 1, *n*) of *f*(*t*), *t* > 0 is defined by

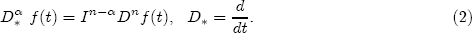

The following properties are some of the main ones of the fractional derivatives and integrals (see [6]-[8], [10], [12], [20], [21]).

Let *β, γ* ϵ *R*^+^ and *α* ϵ (0,1). Then

(i) 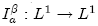, and *f*(*y*) ϵ *L*^1^, then 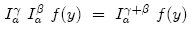.
(ii) 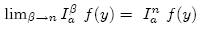 uniformly on [a, b], *n* = 1, 2, 3,…, where 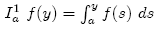.
(iii) 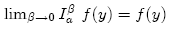 weakly.
(iv) If *f*(*y*) is absolutely continuous on [*a*, *b*], then 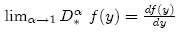.
(v) If *f*(*y*) = *k* ≠ 0, k is a constant, then 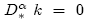.

The following lemma can be easily proved (see [10]).

### Lemma 1

Let *β* ϵ (0,1) if *f* ϵ *C*[0, *T*], then *I*^*β*^ *f*(*t*)|_*t*=0_ = 0.

### 2.1 Equilibrium points and their asymptotic stability

Let *α* ϵ (0, 1] and consider the system ([1]-[3], [11], [13])

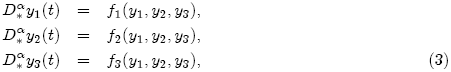

with the initial values

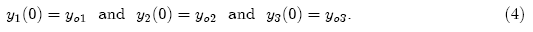

To evaluate the equilibrium points, let

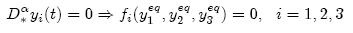

from which we can get the equilibrium points 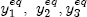.

To evaluate the asymptotic stability, let

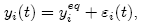

So the the equilibrium point 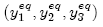 is locally asymptotically stable if the eigenvalues of the Jacobian matrix *A*

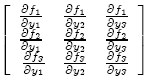

evaluated at the equilibrium point satisfiesis (|arg(*λ*_1_)| > *απ*/2, |arg(*λ*_2_)| > *απ*/2, |arg(*λ*_3_)| > *απ*/2) ([2], [3], [13], [19]). The stability region of the fractional-order system with order *a* is illustrated in Fig. 1 (in which *σ, ω* refer to the real and imaginary parts of the eigenvalues, respectively, and 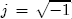). From Fig. 1, it is easy to show that the stability region of the fractional-order case is greater than the stability region of the integer-order case.

**Fig. 1.**
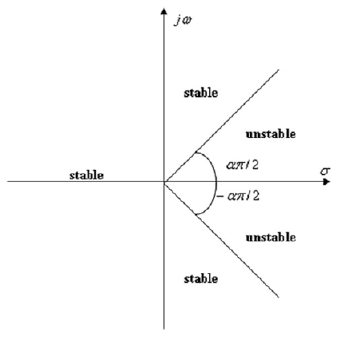
Stability region of the fractional-order system.

The eigenvalues equation of the equilibrium point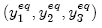 is given by the following polynomial:

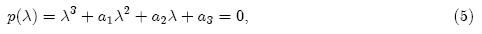

and its discriminant *D*(*P*) is given as:

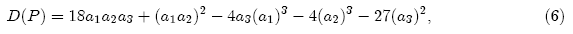

using the results of Ref. [2], we have the following fractional Routh-Hurwitz conditions:

(i) If *D*(*P*) > 0, then the necessary and sufficient condition for the equilibrium point 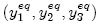, to be locally asymptotically stable, is *a*_1_ > 0, *a*_3_ > 0, *a*_1_*a*_2_ − *a*_3_ > 0.
(ii) If *D*(*P*) < 0, *a*_1_ < 0, *a*_2_ > 0, *α* > 2/3, then all roots of Eq. (5) satisfy the condition |arg(*λ*)| < *απ*/2.
(iii) If *D*(*P*) < 0, *a*_1_ < 0, *a*_2_ < 0, *a*_1_*a*_2_ − *a*_3_ = 0, then 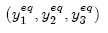 is locally asymptotically stable for *α* ϵ (0, 1).
(iv) The necessary condition for the equilibrium point 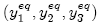, to be locally asymptotically stable, is *a*_3_ > 0.

## 3 Fractional-order SIRS epidemic model on homogenous networks

Let *S*(*t*) be the number of individuals in the susceptible class at time *t*, *I*(*t*) be the number of individuals who are infectious at time *t* and *R*(*t*) be the recovered or vaccinated individuals at time *t* [22].

The fractional-order SIRS epidemic model on homogenous networks is given by

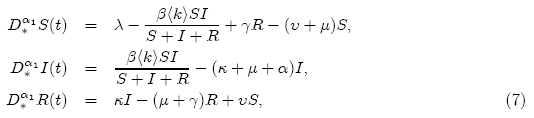

where 0 < *α*_1_ ≤ 1 and the parameters *λ*, *μ*, *β*, *κ*, *υ*, *γ* and α are positive constants, and 〈*k*〉 is the average connectivity in the network neglecting the heterogeneity of the node degrees [22].

Which, together with *N* = *S* + *I* + *R*, implies

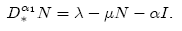

Thus the total population size *N* may vary in time.

To evaluate the equilibrium points, let

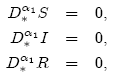

then 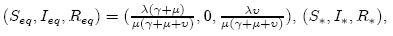, are the equilibrium points where,

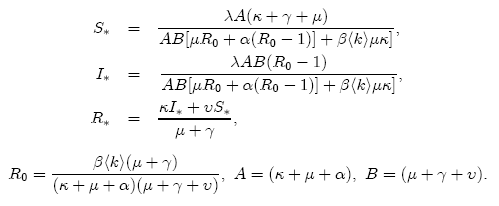

For a disease-free equilibrium point 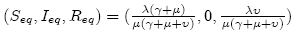 we find that

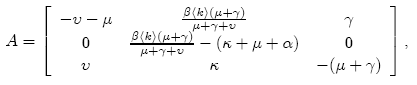

and its eigenvalues are

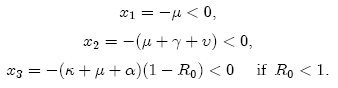

Hence a disease-free equilibrium point 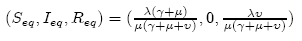 is locally asymptotically stable if *R*_0_ < 1.

For a unique endemic equilibrium point 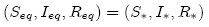 the characteristic polynomial is given by:

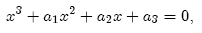

where

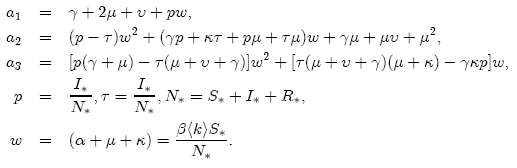

A sufficient condition for the local asymptotic stability of a unique endemic equilibrium point (*S*_eq_, *I_eq_*, *R_eq_*) = (*S*_*_, *I*_*_, *R*_*_) is

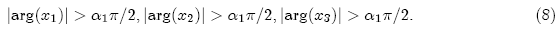

### 3.1 Numerical methods and results

An Adams-type predictor-corrector method has been introduced and investigated further in ([1]”[3], [4], [5], [9]). In this paper we use an Adams-type predictor-corrector method for the numerical solution of fractional integral equations.

The key to the derivation of the method is to replace the original problem (7) by an equivalent fractional integral equations

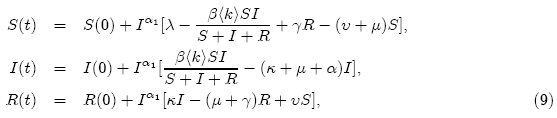

and then apply the **PECE** (Predict, Evaluate, Correct, Evaluate) method.

The approximate solutions displayed in Figs. 2-7 for 〈*k*〉 = 6, *λ* = 1, *β* = 0.2, *μ*, = 0.001, *γ* = 0.1, *α* = 0.00087, *κ* = 0.5 and different 0 < *α*_1_ ≤ 1.In Figs.2-4 we take *υ* = 0.3,5(0) = 450, *I*(0) = 550, *R*(0) = 0 and found that a disease-free equilibrium point 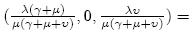 (251.87,0, 748.13) is locally asymptotically stable even if for a large fraction of the infected individuals at the initial time, the disease will eventually disappear where *R*_0_ = 0.602236 < 1.

**Fig. 2.**
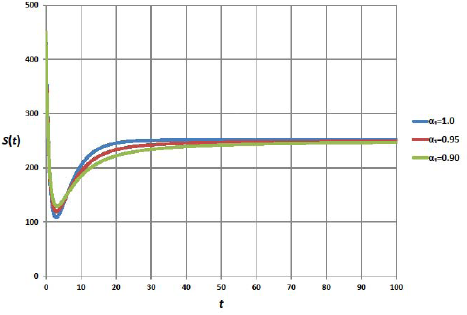

**Fig. 3.**
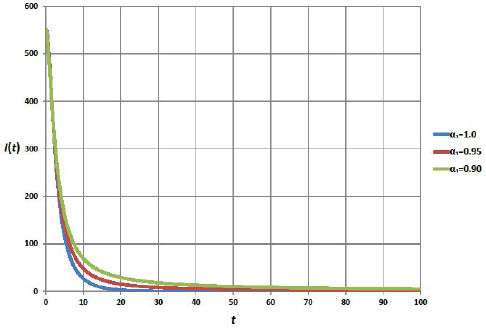

**Fig. 4.**
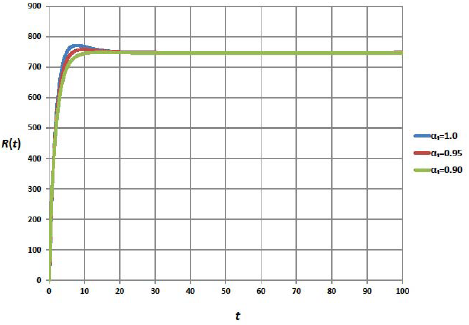

**Fig. 5.**
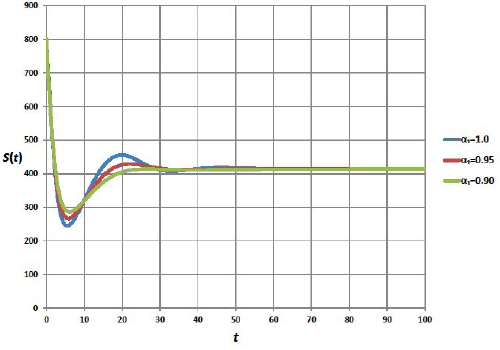

**Fig. 6.**
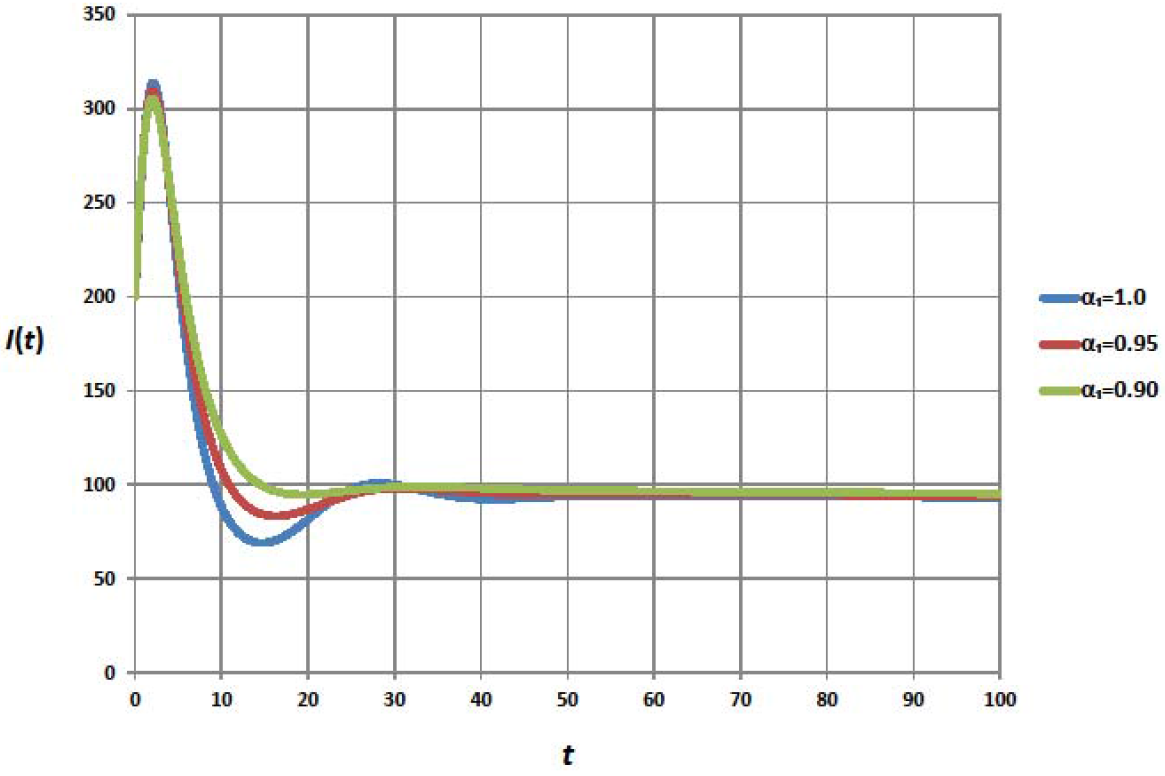

**Fig. 7.**
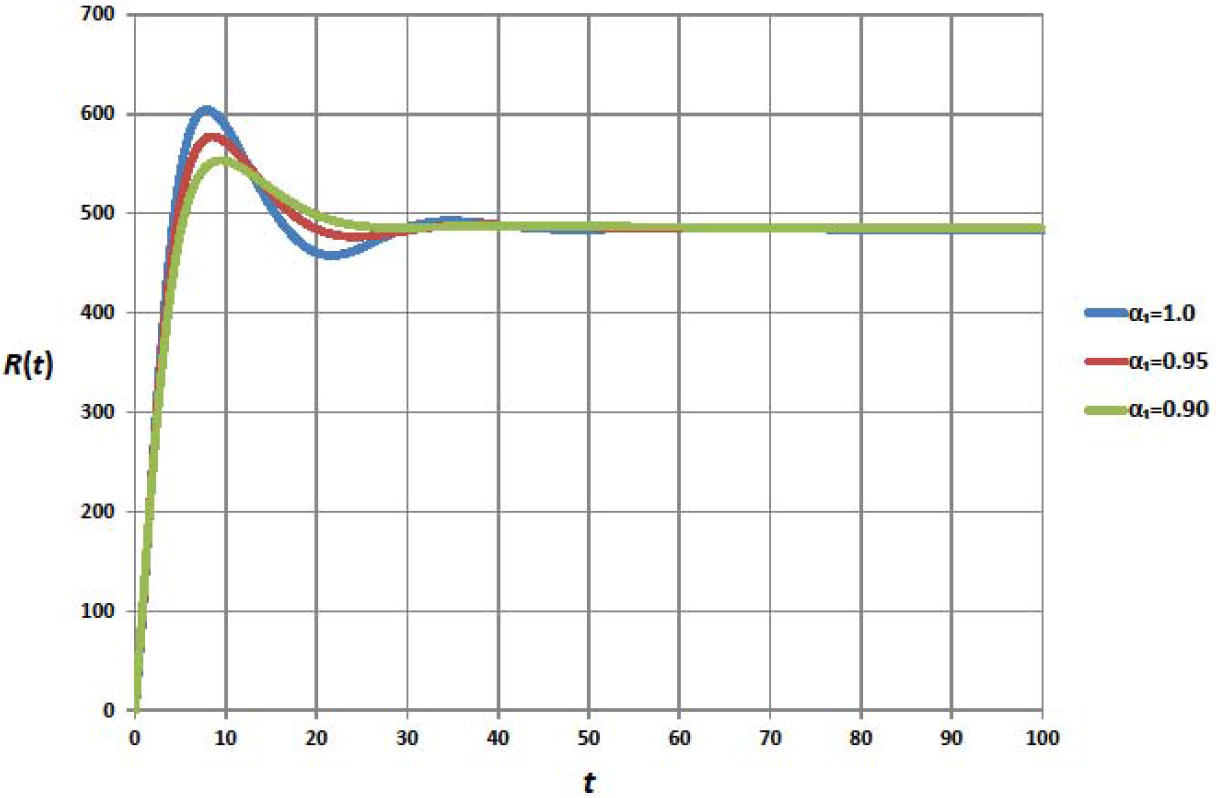

In Figs. 5-7 we take *υ* = 0.005, *S*(0) = 800, *I*(0) = 200, *R*(0) = 0 and found that a disease-free equilibrium point 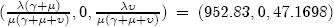 is unstable where *R*_0_ = 2.27827 > 1 and a unique endemic equilibrium point (*S*_*_, *I*_*_, *R*_*_) = (386.518, 87.1414, 450.528) is locally asymptotically stable even if for a small fraction of the infected individuals at the initial time where the condition (8) is satisfied and the eigenvalues are given as:

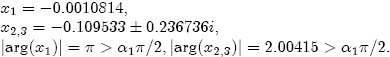

## 4 Conclusions

i- Zika is a fast spreading epidemic.
ii- It is dangerous on pregnant women otherwise it is mild or asymptomatic.
iii- So far it is known to have two transmission routes one via mosquito and the other is via sexual contact.

Therefore we expect that in the future the second route will be more difficult to control. Moreover the fact that Summer Olympics is expected in Brazil in 2016 makes it important to study the epidemic quickly.

Our model will be useful as a conceptual tool for modeling the impact of interventions aiming to control the disease.

## References

[1] E. Ahmed, A. M. A. El-Sayed, E. M. El-Mesiry and H. A. A. El-Saka, Numerical solution for the fractional replicator equation, IJMPC, 16, 7, 1–9, 2005.

[2] E. Ahmed, A. M. A. El-Sayed, H. A. A. El-Saka, On some Routh-Hurwitz conditions for fractional order differential equations and their applications in Lorenz, R¨ossler, Chua and Chen systems, Physics Letters A, 358, 1, 2006.

[3] E. Ahmed, A. M. A. El-Sayed, H. A. A. El-Saka, Equilibrium points, stability and numerical solutions of fractional-order predator-prey and rabies models, J. Math. Anal. Appl., 325, 542–553, 2007.

[4] Kai Diethelm, The Analysis of Fractional Differential Equations: An Application-Oriented Exposition Using Differential Operators of Caputo Type (Lecture Notes in Mathematics), Springer-Verlag Berlin Heidelberg 2010.

[5] E. M. El-Mesiry, A. M. A. El-Sayed and H. A. A. El-Saka, Numerical methods for multi-term fractional (arbitrary) orders differential equations, Appl. Math. and Comput., 160, 3, 683–699, 2005.

[6] A. M. A. El-Sayed, fractional differential-difference equations, Journal of Fractional Calculus, 10, 101–106, 1996.

[7] A. M. A. El-Sayed, Nonlinear functional differential equations of arbitrary orders, Nonlinear Analysis: Theory, Methods and Applications, 33, 2, 181–186, 1998.

[8] A. M. A. El-Sayed and F. M. Gaafar, Fractional order differential equations with memory and fractional-order relayation-oscillation model, (PU.M.A) Pure Math. and Appl., 12, 2001.

[9] A. M. A. El-Sayed, E. M. El-Mesiry and H. A. A. El-Saka, Numerical solution for multi-term fractional (arbitrary) orders differential equations, Comput. and Appl. Math., 23, 1, 33–54, 2004.

[10] A. M. A. El-Sayed, F. M. Gaafar and H. H. Hashem, On the mayimal and minimal solutions of arbitrary orders nonlinear functional integral and differential equations, Math. Sci. Res. J., 8, 11, 336–348, 2004.

[11] A. M. A. El-Sayed, E. M. El-Mesiry and H. A. A. El-Saka, On the fractional-order logistic equation, AML, 20, 7, 817–823, 2007.

[12] R. Gorenflo and F. Mainardi, Fractional Calculus: Integral and Differential Equations of Fractional Order, in A. Carpinteri and F. Mainardi (Eds), Fractals and Fractional Calculus in Continuum Mechanics, Springer, 223–276, 1997.

[13] H. A. El-Saka, E. Ahmed, M. I. Shehata and A. M. A. El-Sayed, On stability, persistence and Hopf Bifurcation in fractional order dynamical systems, Nonlinear Dyn, 56, 121–126, 2009.

[14] H. A. El-Saka, The fractional-order SIR and SIRS epidemic models with variable population size, Math. Sci. Lett. 2, No.3, 1–6, 2013.

[15] H. A. El-Saka and A. El-Sayed, Fractional Order Equations and Dynamical Systems, Lambrt Academic Publishing, Germany (2013), ISBN 978-3-659-40197-8.

[16] H. A. El-Saka, The fractional-order SIS epidemic model with variable population size, Journal of the Egyptian Mathematical Society, 22, pp. 50–54, 2014.

[17] H. A. El-Saka, fractional-order partial differential equation for predator-prey, Journal of Fractional Calculus and Applications, Vol. 5(2) pp. 44–51, July 2014.

[18] H. A. El-Saka, Backward bifurcations in fractional-order vaccination models, Journal of the Egyptian Mathematical Society, Journal of the Egyptian Mathematical Society, 23, 49—55, 2015.

[19] D. Matignon, Stability results for fractional differential equations with applications to control processing, Computational Eng. in Sys. Appl. Vol. 2 Lille France 963, 1996.

[20] I. Podlubny and A. M. A. El-Sayed, On two definitions of fractional calculus, Solvak Academy of science-institute of eyperimental phys. UEF-03–96 ISBN 80-7099-252-2, 1996.

[21] I. Podlubny, Fractional differential equations, Academic Press, 1999.

[22] Yao Hua, Lequan Minab and Yang Kuangc, Modeling the dynamics of epidemic spreading on homogenous and heterogeneous networks, Applicable Analysis: An International Journal, Volume 94, Issue 11, 2015.

